# HIV-1 accessory protein Vpr possesses a cryptic p300-dependent transcription-promoting activity that is blocked by histone deacetylases in CD4^+^ T cells

**DOI:** 10.1101/2025.03.27.645672

**Authors:** Catherine A. Lewis, David M. Margolis, Edward P. Browne

## Abstract

Antiretroviral therapy (ART) has dramatically improved the clinical prognosis for people with HIV and prevents HIV transmission. However, ART does not cure HIV infection because of a persistent, latent reservoir in long-lived cells such as central memory CD4^+^ T (T_CM_) cells. Eliminating or preventing reservoir formation will require a better understanding of HIV-1 latency establishment. We and others have recently shown that host cell factors such as histone deacetylases (HDACs) are critical cellular factors that allow HIV-1 entry into latency. Whether HDACs interact with specific viral factors to regulate latency establishment, however, is unknown. To examine the role of individual HIV-1 accessory proteins, we constructed a panel of HIV-1 reporter strains, each expressing a single HIV-1 accessory protein, and examined them in a primary CD4^+^ T-cell latency model. Interestingly, we found that the HDAC inhibitor (HDACi) vorinostat potently enhances the effect of the HIV-1 protein Vpr in promoting HIV expression in infected cells, suggesting that Vpr possesses a cryptic transcription-promoting activity that is restricted by HDACs. This activity was dependent on a p300-binding domain of Vpr and inhibited by a selective p300 histone acetyltransferase inhibitor. Interestingly, Vpr expression also resulted in a significant increase in the proportion of infected cells with a central memory (T_CM_) phenotype. Furthermore, we observed that T_CM_ cells were more resistant to Vpr-induced apoptosis/cell death than other CD4^+^ T-cell subtypes, indicating that Vpr expression during reservoir formation selects for latent proviruses in T_CM_ cells. Overall, these findings suggest that Vpr plays an important role in shaping the latent reservoir and that HIV-1 latency results, in part, from an HDAC-mediated restriction on Vpr’s transcription-promoting activity. Understanding how viral factors shape the latent reservoir and how host and viral factors interact during HIV-1 latency establishment in CD4^+^ T cells will aid in the development of new latency-targeting therapies.

**Author Summary:** Although antiretroviral therapy is effective at treating HIV, a cure remains elusive. The primary obstacle to HIV cure is the presence of a long-lived reservoir of latently infected cells in which the virus persists despite therapy. Recent work has shown that a sizable fraction of this latent reservoir forms near the time that therapy is initiated, suggesting it may be possible to prevent some of the reservoir from forming. However, latency prevention will require a better understanding of how HIV enters latency, including how viral gene expression is silenced. We therefore sought to examine the role of the interaction between viral proteins and host factors in turning off viral gene expression and found that, whereas the HIV protein Vpr turns on viral gene expression, host histone deacetylases block this activity. Second, we observed that Vpr expression in infected cells leads to an increase in the relative proportion of central memory CD4^+^ T cells, a cell type that harbors latent virus. Our findings on the role of the viral protein Vpr in the silencing of viral gene expression and the persistence of certain memory cell types during infection will be important for developing new approaches to targeting latently infected cells.

## Introduction

Although combined antiretroviral therapy (ART) has significantly reduced human immunodeficiency virus 1 (HIV-1) transmission and has improved prognosis for people with HIV-1 (PWH), ART is still associated with comorbidities (1,2) and PWH still experience problems of access and stigma (3–5). Furthermore, ART is not a cure for HIV infection. ART suppresses viremia by targeting actively replicating virus, but the virus persists as a latent reservoir of stably integrated proviruses and the rate of reservoir decay is too slow for the virus to be cleared during a PWH’s lifetime (6–8). Because of the HIV-1 reservoir, viral levels rapidly rebound if ART is interrupted, demonstrating the urgent need for a cure that either eradicates or permanently silences the latent reservoir.

The latent reservoir begins forming soon after initial infection, and infected cells can undergo clonal expansion to replenish the reservoir even during ART (9–12). One major approach to HIV-1 cure, therefore, has been to develop latency-reversal agents (LRAs) that target host factors involved in HIV-1 transcriptional regulation, with the goal of reactivating viral gene expression to a sufficient extent that the host immune system can detect and kill infected cells. However, while many LRAs can reactivate detectable viral gene expression, latency reversal alone has not yet led to a significant reduction in the frequency of latent infection (13–18). This lack of reservoir reduction is likely due to latency being maintained by multiple levels of epigenetic and transcriptional repression, making it challenging to broadly reactivate the reservoir by modulating a single mechanism.

In recent years, several studies have provided evidence that a large fraction of the long-lived latent reservoir is formed or stabilized near the time of ART initiation (19,20). This finding suggests that it may be possible to intervene during the initiation of ART and prevent a significant part of the reservoir from forming. Because repression of HIV-1 is established gradually in a step-wise manner (21,22), such an approach may be more effective than reversing latency in proviruses after several years of therapy. However, preventing latency will require an improved understanding of how viral latency is established. We previously reported that epigenetic reprogramming via histone deacetylase (HDAC) activity is a critical early step for entry into latency that licenses subsequent repressive histone methylation (23). Furthermore, we showed that HDAC activity may also help to maintain T cells in a long-lived, stem cell memory-like state (T_SCM_ cells), a memory subset in which the long-lived latent reservoir has been reported to persist (23,24).

In this study, we sought to understand how HIV-1 accessory proteins shape viral entry into latency, both alone and in combination with HDACs. We report that the viral accessory protein Vpr has a strong effect that appears to be more than additive on viral gene expression when combined with the class I HDAC inhibitor (HDACi) vorinostat, suggesting that Vpr possesses a cryptic activity in CD4^+^ T cells that counteracts latency establishment but is blocked by HDACs. Furthermore, this activity depends on a p300-binding domain of Vpr and histone acetyltransferase activity of p300. We also observed that Vpr has a striking effect on the phenotype of cells that enter the pool of latently infected cells by selecting for cells with a long-lived T_CM_ phenotype. Our data suggest that Vpr expression during latency establishment plays an important role in shaping key characteristics of the latent reservoir.

## Results

### Vpr expression antagonizes HIV latency establishment in the presence of a histone deacetylase inhibitor (HDACi)

To investigate the role of HIV-1 accessory proteins in the latency establishment, we used a primary CD4^+^ T-cell model of HIV latency that we have previously established. This model uses a reporter strain of HIV (HIV-GFP) that lacks expression of all viral proteins except Tat and Rev (21,25,26) but expresses a destabilized eGFP gene and the surface marker Thy1.2. In this model, activated CD4^+^ T cells are infected with HIV-GFP before being cultured for up to three weeks. Over this three-week period in this single-round infection system, viral gene expression progressively declines and a latently infected population with low to undetectable GFP expression emerges from the actively infected population. To study the effect of individual HIV-1 proteins on latency initiation in this system, we generated single gene revertant clones of HIV-GFP for *Gag/Pol*, *Vif*, *Vpu*, *Vpr*, and *Nef*. We confirmed successful reversion of each gene and rescue of Gag/Pol, Vif, and Vpr protein production by sequencing of the plasmid and western blot of transfected 293T cells, respectively (Fig S1A). Expression of the two reporter genes, Thy1.2 and GFP, was confirmed in infected CD4^+^ T cells for all revertant viruses (Fig S1D). Of note, CD4 expression was downregulated in cells infected with Nef or Vpu revertant virus, confirming functional rescue of these HIV-1 proteins (Fig S1B).

We then examined the effect of each viral gene individually on establishment of HIV latency in CD4^+^ T cells. Recently activated CD4^+^ T cells were infected with infectious supernatant for HIV-GFP as well as each of the revertant virus clones, and, over the following three weeks, flow cytometry was used to measure the percentage of productively infected cells (%GFP^+^ cells within the Thy1.2^+^ population) and the median fluorescence intensity (MFI) of GFP expression within the GFP^+^ population. When we examined the overall effect of the revertant viruses, we did not observe a large difference between the revertants and the parental virus strain for Gag/Pol, Nef, Vif or Vpu. Although previous studies have shown that Vpr can transactivate HIV-1 expression (27–32), we also did not observe a strong effect of Vpr or any of the other reverted viral proteins on the percentage productively infected cells in this model system and only a slight positive effect of Vpr on viral gene expression by GFP median fluorescence intensity (MFI; Fig 1D,F). These results indicate that, individually, the HIV proteins Gag/Pol, Vif, Vpr, and Nef do not have a strong effect on HIV transcriptional downregulation or latency in infected CD4^+^ T cells.

**Fig 1:**
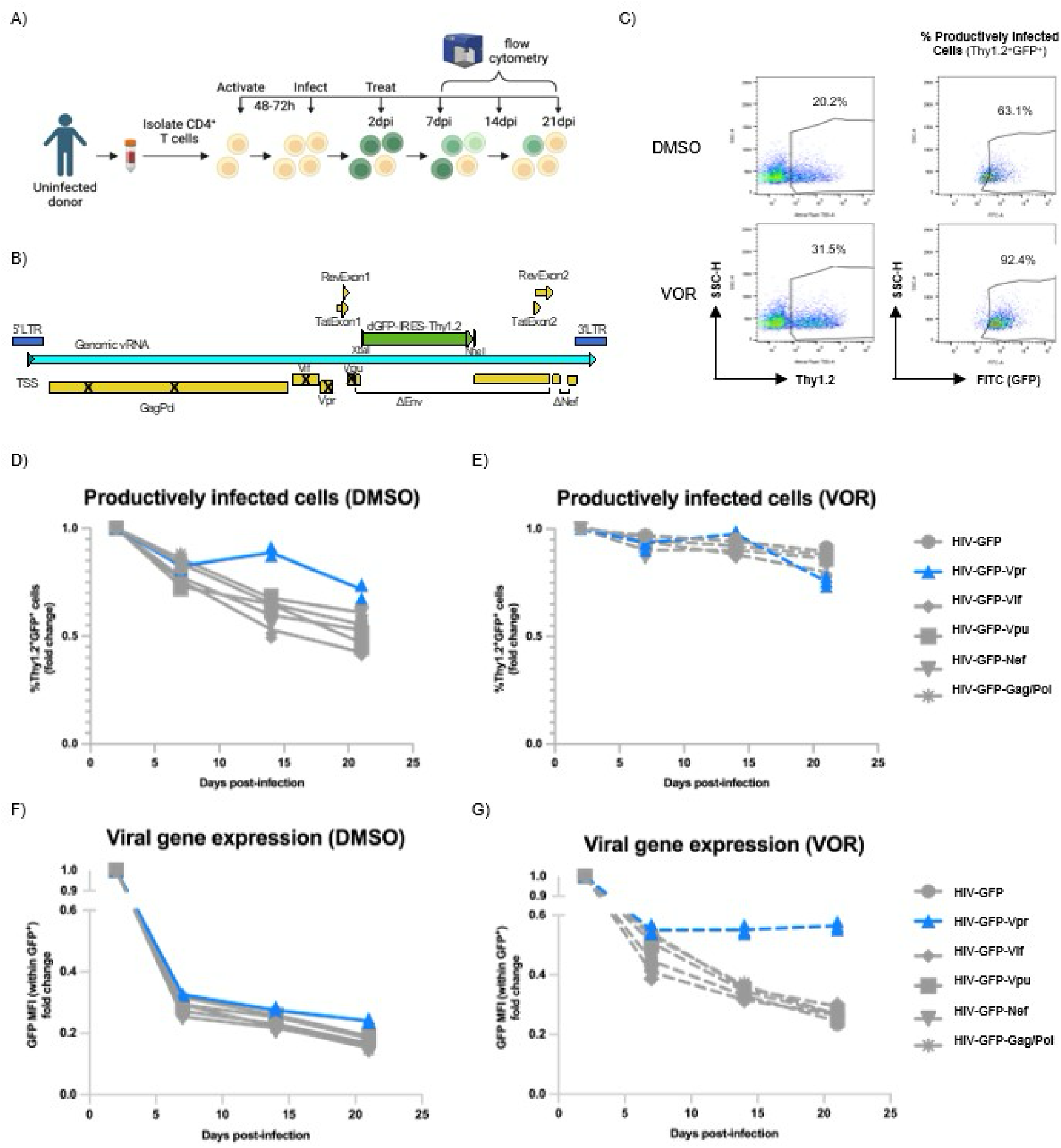
Effect of single viral gene revertants on productive infection and viral gene expression. A-B) Primary CD4^+^ T-cell model A) Overall workflow (created with Biorender.com). B) HIV-GFP-Thy1.2 construct. Bold Xs represent point mutations that result in premature stop codons. ΔEnv represents the portion of the Env-encoding region that has been replaced with a destabilized eGFP followed by an IRES-Thy1.2 cassette. ΔNef indicates a truncation in the Nef-encoding region. TSS = transcription start site. C) Gating strategy for Thy1.2, GFP, and representative plot of median fluorescence intensity (MFI) within GFP+ cells. Top plots are vehicle control (DMSO)-treated. Lower plots are vorinostat (VOR)-treated. D-E) %productively infected cells (Thy1.2^+^GFP^+^) following infection with revertant virus and D) DMSO or E) VOR treatment. F-G) MFI within productively infected cells following F) DMSO or G) VOR treatment. n = 2 for all experiments.

We previously showed that, in this system, histone deacetylases (HDACs) play an important role in initiating transcriptional silencing of the HIV-1 promoter (long terminal repeat; LTR) during latency establishment (21). We therefore sought to assess the relationship between individual viral proteins and HDACs by treating cells with the class I HDAC inhibitor (HDACi) vorinostat or vehicle control (DMSO) from two days post-infection onwards. As we previously observed, vorinostat alone resulted in a large fraction of the infected cells failing to enter latency and remaining GFP^+^, whereas a substantial fraction of DMSO control-treated cells became GFP^-^ by 21dpi. Interestingly, although we did not see a significant change in the percentage of productively infected (GFP^+^) cells following infection with any of the revertant viruses (Fig 1E), vorinostat exposure led to a significant increase in viral gene expression (GFP MFI within the GFP^+^ cell gate) in cells infected with Vpr-expressing revertant compared with cells infected with parental virus HIV-GFP (Fig 1G, Fig 2C, data with statistical analysis from 9 donors combined in Fig S2A). The combination of vorinostat treatment and Vpr expression led to strongly maintained viral gene expression over time, an effect that was not observed with either vorinostat or Vpr alone (Fig 1D,G; Fig 2C). Overall, these data indicate that Vpr possesses an activity that can potently counteract downregulation of viral gene expression in CD4^+^ T cells but is blocked by the action of a histone deacetylase.

**Fig 2:**
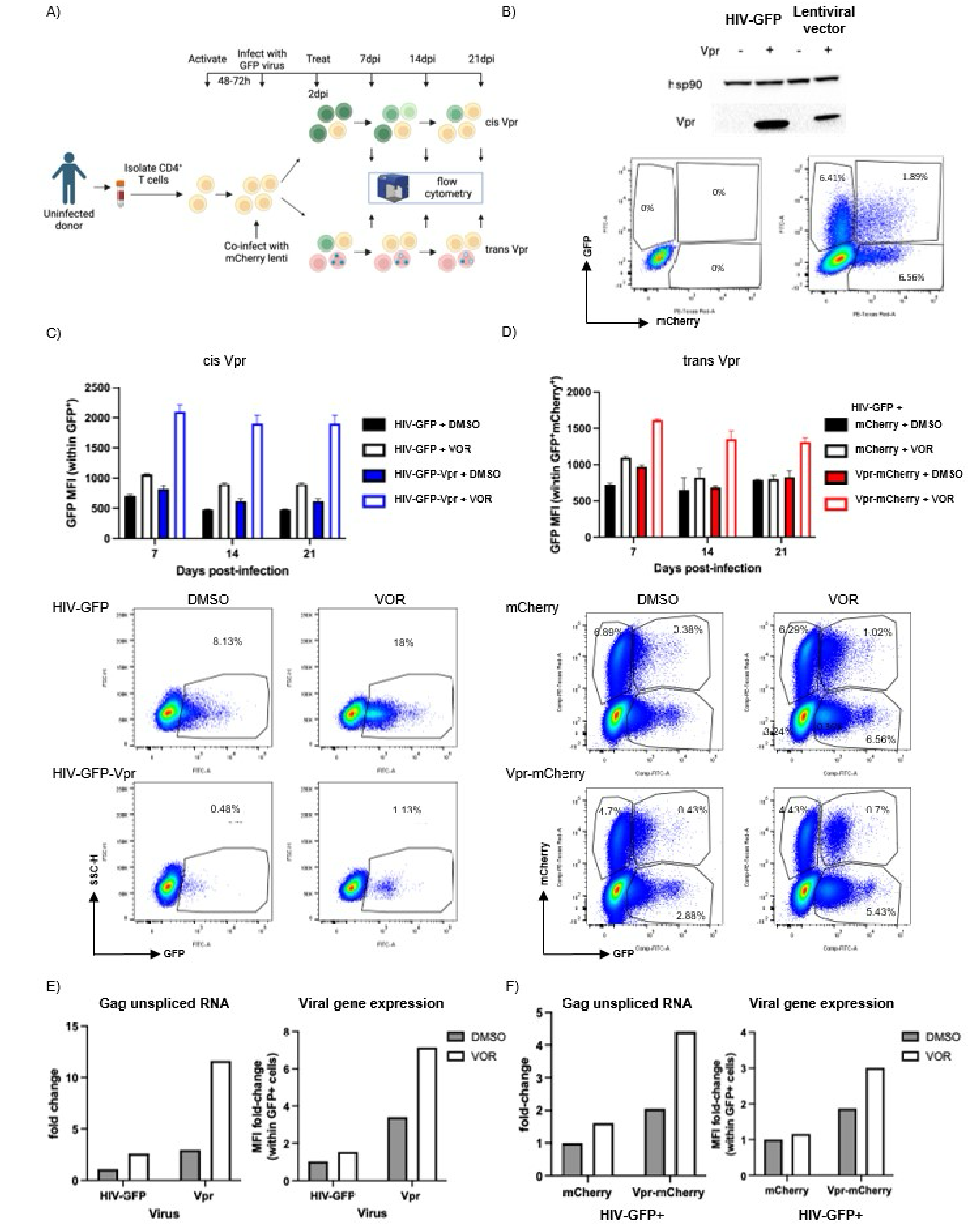
Viral gene expression within productively infected (GFP^+^) cells is increased in presence of Vpr and vorinostat (VOR). A) Top panel (cis Vpr expression): schematic of workflow with infection with Vpr revertant vs. Vpr-null virus (HIV-GFP) and lower panel (trans Vpr expression): co-infection with Vpr-mCherry lentivirus or mCherry lentivirus and HIV-GFP. B) Top panel: western blot for Vpr expression in 293T cells transfected with HIV-GFP, Vpr revertant plasmid, mCherry lentivirus plasmid, or Vpr-mCherry lentivrus plasmid. Lower panel: representative dot plot of co-infected CD4^+^ T cells. C-D) Representative quantification of GFP MFI over 21d in C) cis Vpr infection model (n = 9) and D) trans Vpr infection model (n = 6). Representative dot plots with and without VOR below. E-F) Left: Representative GFP MFI quantification by flow cytometry and (right) gag unspliced transcript quantification by qPCR in E) cis Vpr infection model (n=2) and F) trans Vpr infection model (n=3).

### Vpr expression *in cis* or *in trans* in combination with an HDACi enhances HIV expression at the transcriptional level

Because the HIV LTR is repressed by HDACs during latency and the *Vpr* gene of HIV is expressed in an LTR-dependent manner, we hypothesized that the strong combined effect of HDACis and Vpr expression might be due to upregulation of Vpr expression itself by HDACis. Thus, to determine whether Vpr in combination with HDACi enhances HIV expression when Vpr expression is LTR-independent, we used a co-infection model: Primary CD4^+^ T cells were co-infected with HIV-GFP and an EF1*α*-driven lentiviral vector expressing either mCherry alone or co-expressing Vpr and mCherry (mCherry and Vpr-mCherry, respectively) followed by vorinostat treatment beginning at 2-3dpi and monitored by flow cytometry for 21 days ( Fig 2A,B). Similar to our results with cis Vpr expression, co-infected cells (GFP^+^mCherry^+^) with trans Vpr expression and treated with vorinostat maintained viral gene expression to a greater extent than either vorinostat-treated cells co-infected with mCherry lentivirus or DMSO-treated cells co-infected with the Vpr-mCherry lentivirus (Fig 2D, data with statistical analysis from 6 donors combined in Fig S2B). Together, these results indicate that HDACs block a transactivation activity of Vpr in CD4^+^ T cells, whether Vpr is expressed in cis or trans, and that the potent combined effect of Vpr and vorinostat is independent of the promoter driving Vpr expression.

Because viral gene expression is measured via levels of a reporter protein (GFP), these data could reflect an effect on viral transcription or also an effect on a post-transcriptional step such as RNA splicing, translation, or protein stability. To determine whether HIV-1 transcription levels are increased following Vpr expression and vorinostat treatment, at 14dpi, we sorted GFP^+^ cells from vorinostat or DMSO-exposed cells (n = 2) that were infected with HIV-GFP-Vpr or HIV-GFP and measured Gag unspliced mRNA (Gag usRNA) by real-time qPCR. We observed a 13-fold increase in Gag usRNA with combined HIV-GFP-Vpr infection and vorinostat exposure compared with HIV-GFP infection with DMSO exposure. By contrast, we observed approximately 2.5-fold increases in Gag usRNA with either HIV-GFP-Vpr infection or vorinostat treatment alone (Fig 2E, Fig S2C). We also confirmed this observation with lentiviral-driven Vpr expression in trans (Fig 2F, Fig S2D, n = 3). Overall, these data demonstrate that Vpr and vorinostat combine to significantly upregulate HIV expression and counteract latency establishment at the transcriptional level.

### Vorinostat treatment does not alter Vpr’s sub-cellular localization

Next, we hypothesized that vorinostat could modify or enhance Vpr’s transcription-enhancing activity by affecting its subcellular localization. In previous studies, largely performed in cell lines, Vpr has primarily been found in the nucleus and nuclear envelope but may also be present in the cytoplasm and could affect HIV expression through its activity in either compartment (33–35). For example, vorinostat could potentially promote increased nuclear import of Vpr, thereby allowing it to directly enhance HIV expression in the nucleus. To assess whether vorinostat treatment affects Vpr localization within infected cells, we fractionated vorinostat- or DMSO-treated cells that had been infected with either HIV-GFP or HIV-GFP-Vpr into chromatin, soluble nuclear, membrane, and cytosolic fractions and then examined the abundance of Vpr in each compartment by western blotting for Vpr. To confirm successful fractionation of the cells, we examined the distribution of the cytoplasmic protein beta-tubulin, the soluble nuclear protein Lamin B, and the chromatin bound protein Histone 3 (H3). Vpr was abundantly present in three of the four fractions (cytoplasmic, membrane, and soluble nuclear) at relatively equivalent amounts, while Vpr was not abundantly detected within the chromatin fraction. Notably, vorinostat exposure had no detectable effect on the abundance of Vpr in any sub-cellular compartment, indicating that vorinostat does not enhance the transcription-promoting activity of Vpr by modulating Vpr localization (Fig S3).

### Vpr induces G2/M arrest and apoptosis in primary CD4^+^ T-cell model

Although the function of Vpr in HIV-1 infection and replication in macrophages is relatively well-characterized, the role of Vpr in CD4^+^ T cells is less well understood. Studies have suggested that Vpr induces G2/M arrest and apoptosis in CD4^+^ T cells (36–38), and some evidence suggests that G2/M arrest may be correlated with increased viral gene expression (29,31). We therefore sought to confirm that Vpr expression induces G2/M arrest and apoptosis in our primary CD4^+^ T-cell model. Using DAPI, a DNA fluorophore that stoichiometrically binds dsDNA, thereby allowing resolution of cell cycle phases based on the DNA content of each cell, we observed that cells productively infected with HIV-GFP-Vpr had a significantly higher proportion of cells (∼46%) in G2/M phase than cells productively infected with HIV-GFP (∼17%). this finding suggests that Vpr arrests cell cycle at G2/M in our primary cell infection model (Fig S4A). This effect was not observed in uninfected cells. As previously observed, vorinostat also induced G2/M arrest (39–41) in both HIV-GFP-Vpr- and HIV-GFP-infected cells.

To assess apoptosis, we stained infected cells at two days post-infection with Zombie Violet (ZV; BioLegend), a fluorescent amine-binding viability dye that is excluded from live cells, and Annexin V, which binds phosphatidylserine, a phospholipid that is exposed on the surface of apoptotic cells (42). Notably, we observed elevated levels of both early Annexin V^+^, ZV^-^) and late apoptotic/dead cells (Annexin V^+^, ZV^+^) in samples infected with Vpr-expressing HIV than those infected with HIV-GFP. This result demonstrates that Vpr induces apoptosis and likely other types of cell death in our primary CD4^+^ T-cell model as in other HIV-1 infection models (36,43–45) (Fig S4B).

### Combined effect of Vpr and vorinostat on HIV expression depends on the p300-binding domain of Vpr

In addition to inducing G2/M arrest and apoptosis, Vpr has been shown to interact with numerous cellular factors, notably, DDB1-Cul4-associated factor 1 (DCAF1), which targets host factors for proteasomal degradation. Another cellular factor identified by biochemical assay to interact with Vpr is the histone acetyltransferase (HAT) p300 (28,46), a transcriptional co-activator that modulates transcription via chromatin remodeling and by binding to transcription factors, such as NF-*κ*B. To assess the mechanism by which Vpr activates viral gene expression in the presence of vorinostat, we generated a set of previously characterized Vpr mutants in both the HIV-GFP and lentivirus backgrounds. These Vpr mutants were 1) G2/M arrest induction (Y50A)(47), 2) G2/M arrest and apoptosis induction (R90K)(48), 3) DCAF1 binding (Q65R)(49), and 4) p300 binding (F72A/R73A)(28).

We first confirmed Vpr expression for each of the mutant Vpr-expressing viruses by western blot of transfected 293T cells (Fig S5A,B). Notably, three of the four mutants, F72A/R73A, Y50A, and R90K, were expressed at a lower level when expressed in cis (encoded by HIV-GFP) but not when encoded within the recombinant Vpr expression lentivirus. To assess whether any of the four Vpr mutants exhibited altered activity in our primary CD4^+^ T-cell latency model, we then infected activated CD4^+^ T cells with HIV-GFP, HIV-GFP-Vpr or one of the four HIV-GFP-Vpr mutant viruses and then treated the infected cells with vorinostat or vehicle control (DMSO). When we measured the effect on viral gene expression at 14dpi, we observed that the F72A/R73A mutant exhibited reduced ability to activate HIV gene expression in the presence of vorinostat. By contrast, the Q65R, Y50A and R90K mutants were all still able to induce HIV expression in combination with vorinostat to a similar degree as wild type Vpr (Fig 3A). These data suggest that the p300-binding domain of Vpr is required for HDACi-dependent transactivation of viral gene expression. Although expression of the F72A/R73A was reduced in transfected 293T cells relative to wild type Vpr, the other Vpr mutants with reduced expression (Y50A and R90K) retained full activity for promoting HIV expression when combined with vorinostat, suggesting that this phenomenon is not exquisitely sensitive to Vpr expression levels.

**Fig 3:**
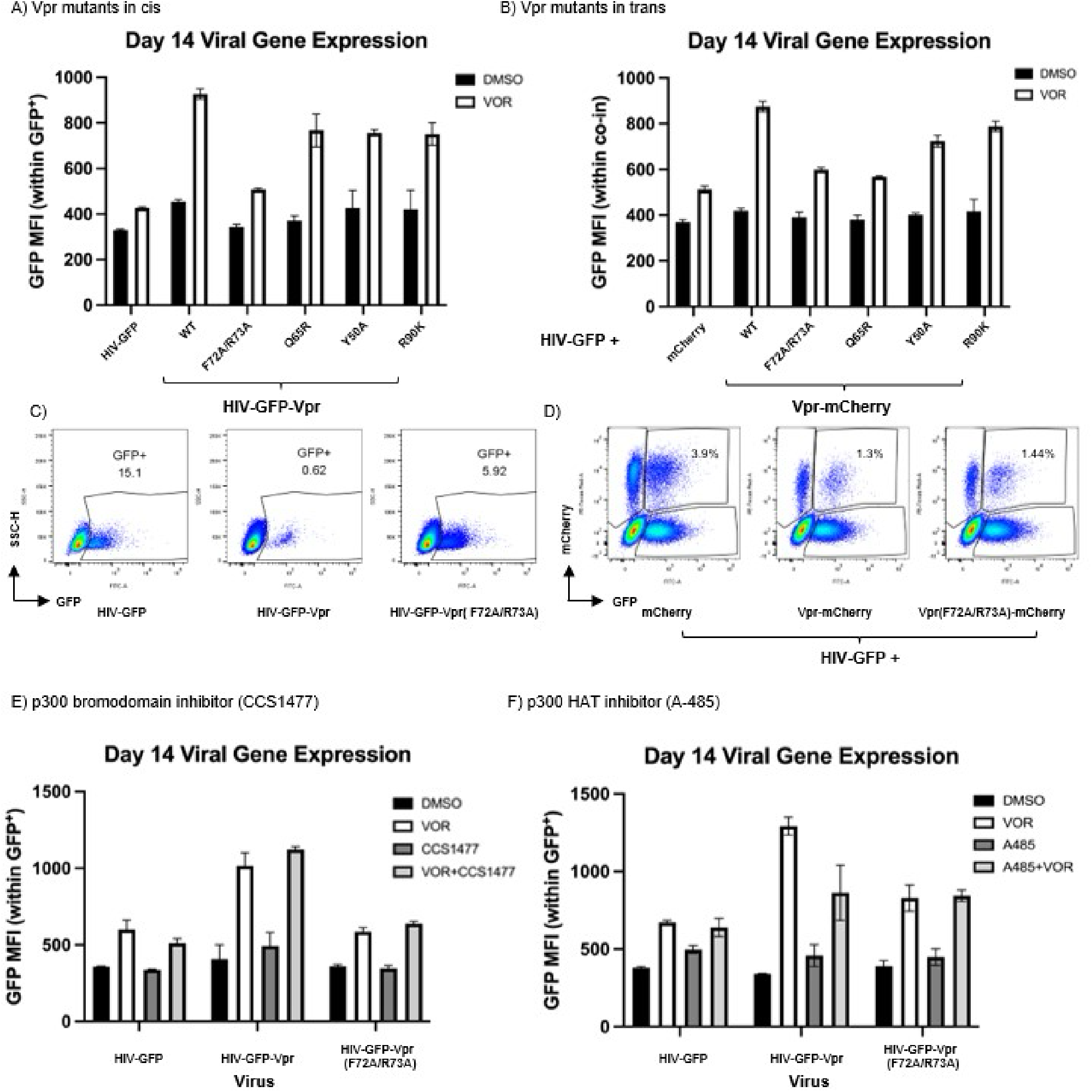
Vpr p300-binding site and p300 acetyltransferrase activity required for synergy between Vpr and vorinostat (VOR). A-B) Day 14 post-infection viral gene expression (GFP median fluorescence intensity; MFI) within DMSO- or VOR-treated CD4^+^ T cells productively infected (GFP^+^) with A) HIV-GFP or HIV-GFP-Vpr mutant virus or B) co-infected with HIV-GFP and mCherry or Vpr-mCherry mutant lentivirus. Representative of two donors with three technical replicates each. C) Representative dot blots from A) for cells infected with HIV-GFP, HIV-GFP-Vpr, or HIV-GFP-Vpr (F72A/R73A) and treated with VOR. D) Representative dot blot from B) for cells co-infected with HIV-GFP and Vpr (F72A/R73A)-mCherry, Vpr-mCherry, or MCherry and treated with VOR. E-F) CD4^+^ T cells infected with HIV-GFP, HIV-GFP-Vpr, or HIV-GFP-Vpr (F72A/R73A) virus and treated with DMSO, VOR, or E) p300 bromodomain inhibitor (CCS1477) alone or in combination with VOR or F) p300 histone acetyltransferrase inhibitor (A485) alone or in combination with VOR. Representative of three donors with three technical replicates each. VOR = vorinostat

When we examined the activity of these Vpr mutants following co-infection with HIV-GFP and lentiviral-driven Vpr expression in trans (Fig 2A), we also observed a reduced ability of the F72A/R73A mutant to drive viral gene expression in combination with vorinostat (Fig 3B). Of note, Vpr (F72A/R73A) was expressed at a similar level to wild-type Vpr in lentivirus-transfected 293T cells (Fig S5B), indicating that the reduced ability of this mutant to promote HIV gene expression was not due to lower Vpr expression. Interestingly, HIV gene expression was also lower in infected cells that were transduced with a lentivirus expressing the Q65R mutant compared with wild-type Vpr (Fig S5B), although this may have been related to the overexpression of Vpr when cells were infected with Vpr (Q65R)-mCherry. Overall, these data indicate that the p300-binding domain of Vpr is required for its ability to promote high HIV-1 expression in the presence of vorinostat.

### A chemical inhibitor targeting the p300 histone acetyltransferase domain inhibits the combined effect of Vpr and vorinostat on HIV-1 expression

Given our observation that the p300-binding domain of Vpr is required for the combined pro-transcriptional effect of Vpr and vorinostat, we next sought to further probe the mechanism of this effect using p300-targeted inhibitors. p300 has five protein-binding domains in addition to a histone or protein acetyltransferase (HAT/PAT) domain and a bromodomain (BRD) that mediates binding to acetylated histones. We first assessed the effect of the p300-specific bromodomain inhibitor CCS1477 (50) on HIV-1 gene expression when Vpr expression is combined with vorinostat treatment. We infected primary CD4^+^ T cells with HIV-GFP, HIV-GFP-Vpr, or HIV-GFP-Vpr (F72A/R73A). Two days post-infection, we exposed the cells to DMSO, vorinostat, p300 bromodomain inhibitor (CCS1477), or a combination of vorinostat and CCS1477 and monitored GFP expression for 14 days. Notably, we did not observe any effect of CCS1477 alone or in combination with vorinostat on the level of viral gene expression in infected cells (measured by GFP MFI; Fig 3E; 3 donors combined in Fig S6A). Next, to assess the role of the histone acetyltransferase domain of p300 in the combined effect of Vpr and vorinostat, we again performed experiments in CD4^+^ T-cells infected with HIV-GFP, HIV-GFP-Vpr, or HIV-GFP-Vpr F72A/R73 but instead exposed the infected cells to a p300 HAT inhibitor (A-485). To select an A-485 concentration, we performed a 6-point, 3-fold dose-response curve from 5-0.02uM and selected 0.56uM as the highest concentration at which cells were still viable. A-485 has been reported to have high selectivity for the p300 HAT domain over other HAT family members and BET bromodomain proteins (51). We observed that the addition of A-485 attenuated the effect of vorinostat and Vpr to a level similar to that seen with the F72A/R73A Vpr mutant that lacks p300-binding activity (Fig 3F; 3 donors combined in Fig S6B). This result suggests that HAT activity of p300 contributes to the combined strong pro-transcriptional effect of vorinostat and Vpr and that the p300 bromodomain is not required for this interaction.

### Combined vorinostat exposure and Vpr expression induces a unique transcriptional signature in HIV infected cells

Having observed that the combination of vorinostat exposure and Vpr expression enhanced HIV-1 expression beyond each individual condition alone, we next hypothesized that combined vorinostat exposure and Vpr expression might also induce a unique effect on the transcriptome of infected cells. To investigate this hypothesis, CD4^+^ T cells from three different seronegative donors were infected with either HIV-GFP or HIV-GFP-Vpr and treated with either DMSO or vorinostat for 12 days. As expected, we observed that the combined effect of vorinostat and Vpr expression potently upregulated HIV expression (Fig 5A). We then isolated RNA from sorted infected (GFP^+^) cells and performed bulk RNA sequencing. When we examined the overall structure of the RNAseq data with principal component analysis (PCA; Fig 5B), we observed that the datapoints could be separated based on condition, indicating a consistent biological signal across donors. We then determined sets of differentially expressed genes (DEGs) using DESeq2. Using an adjusted p-value of 0.05 as the significance cutoff, we observed 177 upregulated and 361 downregulated genes in vorinostat- vs. vehicle control-treated cells infected with HIV-GFP. When comparing cells infected with HIV-GFP-Vpr vs. HIV-GFP, there were 447 upregulated and 367 downregulated genes. Notably, cells treated with vorinostat and infected with HIV-GFP-Vpr exhibited a total of 1388 upregulated vs. 1239 downregulated genes compared with DMSO-exposed cells infected with HIV-GFP (Fig 5C-G). These data indicate that the combined vorinostat/Vpr condition induces a much more potent transcriptional signature in HIV-1-infected CD4^+^ T cells than vorinostat or Vpr alone. Thus, the potent effect of Vpr/vorinostat is also observed with numerous cellular genes and is not specific to HIV. These observations provide further evidence that Vpr has a strong transcriptional effect in CD4^+^ T cells that is blocked by HDACs and revealed by vorinostat treatment.

**Fig 4:**
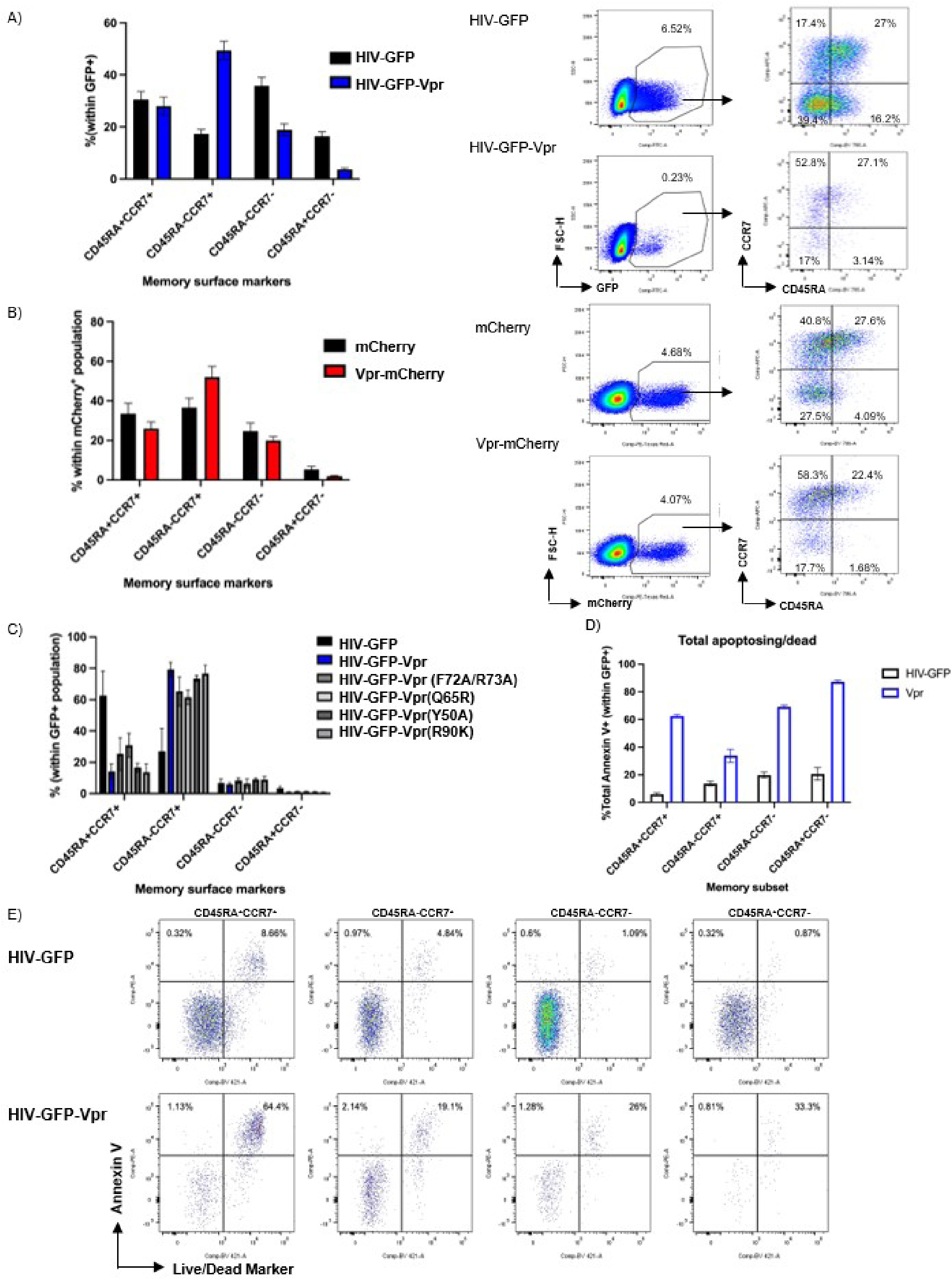
Vpr expression results in increased % cells with T_CM_-like phenotype (CD45RA- CCR7+), potentially because of relatively lower cell death. A-B) Representative quantification of %cells in each of four memory compartments as defined by CD45RA and CCR7 surface marker expression in CD4^+^ T cells infected with A) HIV-GFP or HIV-GFP-Vpr (n = 6) or B) mCherry or Vpr-mCherry lentivirus (n = 5) with representative dot plots C) % cells in each of four compartments following infection with HIV-GFP-Vpr mutant viruses (n =2). D) %Annexin V^+^ cells within GFP^+^ CD4+ T cells in each of four CD4^+^ T-cell memory compartments and E) representative flow plot from each quadrant of CCR7 vs. CD45RA dot plots (n = 5).

**Fig 5:**
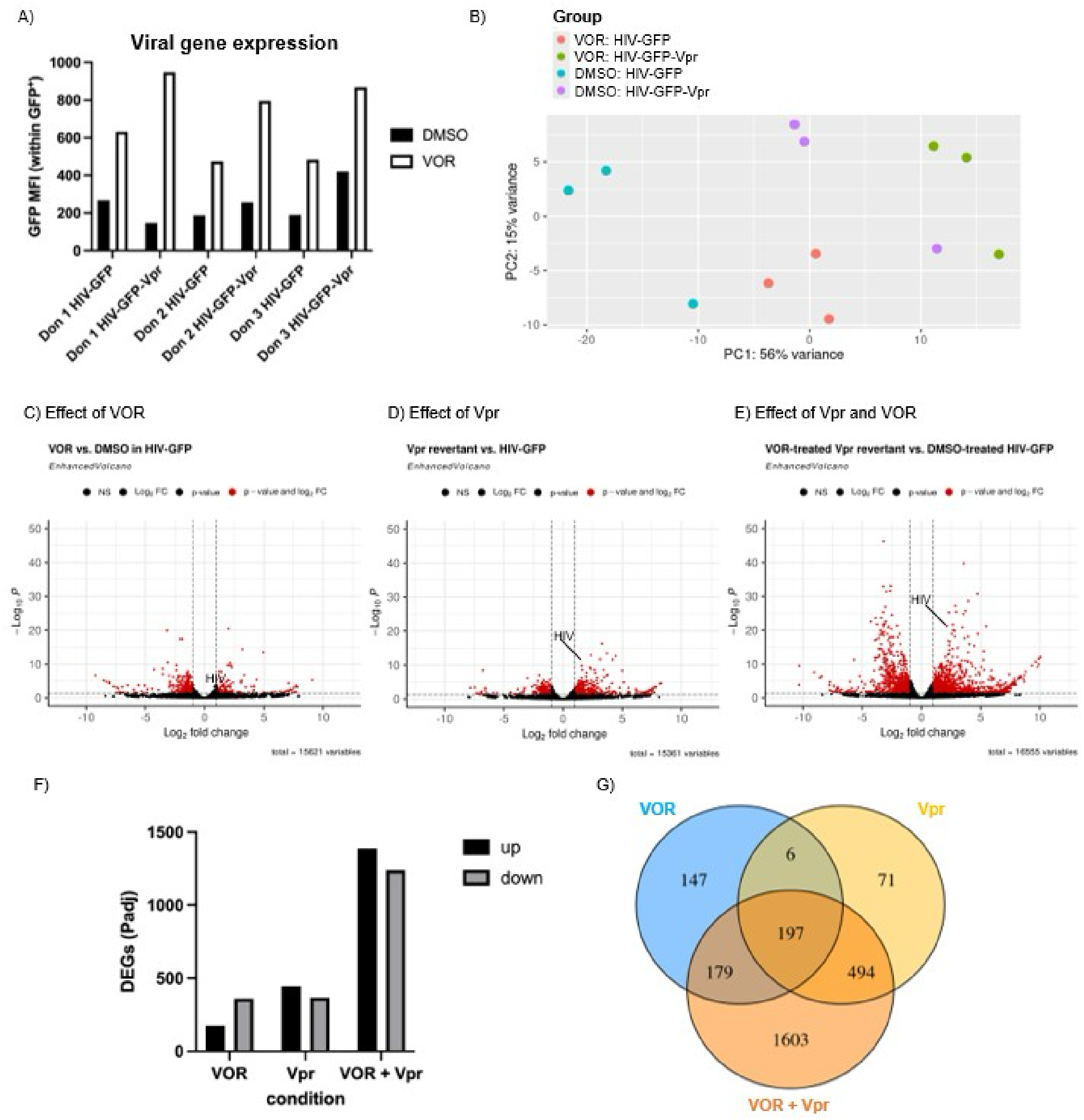
Many more differentially expressed genes (DEGs) result from VOR treatment and Vpr expression than either condition alone. A) Primary CD4^+^ T cells from 3 donors were infected with either HIV-GFP or HIV-GFP-Vpr and treated with 500nM vorinostat starting 2 days post-infection (dpi). 14dpi, cells were analyzed by flow cytometry, and RNA was isolated for bulk RNA sequencing (RNASeq). A) Flow cytometry results for GFP median fluorescence intensity (MFI). B) Principal component analysis of RNASeq samples from three donors. C-E) Volcano plots of DESeq2 results for C) VOR- vs. DMSO-treated cells infected with HIV-GFP, D) DMSO-treated cells infected with HIV-GFP-Vpr or HIV-GFP, or E) VOR-treated, HIV-GFP-Vpr-infected cells vs. DMSO-treated, HIV-GFP-infected cells. Y-axis is based on adjusted P-value. Alpha = 0.05. F) Quantification of DEGs from C-E. G) Venn diagram of DEGs from C-E.

We next examined whether any specific biological pathways were affected by Vpr and vorinostat either alone or in combination. Gene set enrichment analysis (GSEA) using the Hallmark gene set (52) confirmed downregulation of pathways involved in G2/M checkpoint, mitosis, and MTORC1 signaling in the presence of vorinostat, as expected and as we have previously reported (21). We observed a similar effect for Vpr alone (Fig S7F). Interestingly, IL6/Jak/Stat3 signaling was downregulated in vorinostat- vs. vehicle control-treated HIV-GFP-Vpr-infected cells. In addition, we observed an increase in the expression of genes involved in “TNF*α* signaling via NF-*κ*B” in HIV-GFP-Vpr-infected cells treated with vorinostat vs. HIV-GFP-infected cells treated with vorinostat. We did not, however, observe enrichment for any unique pathways in HIV-GFP-Vpr-infected cells treated with vorinostat vs. HIV-GFP-infected cells treated with DMSO. Overall, these data suggest that, while signaling through NF-*κ*B may play a role in the increased HIV-1 transcription in the presence of vorinostat and Vpr, the combined effect of Vpr and vorinostat on host cells is likely broad and involves multiple pathways.

### Vpr expression in HIV-1-infected cells increases the proportion of cells with a central memory phenotype

Given the potent effect of Vpr on the transcriptome of infected cells, we hypothesized that Vpr might affect the T-cell memory subset phenotype of the infected cells. CD4^+^ T cells in vivo can be subdivided into functionally distinct subsets and differentiate along a linear trajectory from naïve (T_N_) cells to central memory cells (T_CM_) to effector memory cells (T_EM_) and finally to terminally differentiated effector memory cells (T_EMRA_). We therefore sought to determine whether Vpr modulates memory-cell composition in activated primary CD4^+^ T cells during latency establishment. Based on the two cell surface markers CCR7 and CD45RA, we defined these four populations of memory CD4^+^ T cells as follows: CD45RA^+^CCR7^+^ stem cell memory (T_SCM_), CD45RA^-^CCR7^+^ central memory T cells (T_CM_), CD45RA^-^CCR7^-^ effector memory T cells (T_EM_), and CD45RA^+^CCR7^-^ effector memory re-expressing RA T cells (T_EMRA_). Interestingly, at weeks 1, 2, and 3 post-infection, we observed an increased fraction of cells with a T_CM_-like phenotype following infection with HIV-GFP-Thy1.2-Vpr compared with HIV-GFP infection (Fig 4A, 6 donors combined in Fig S7A). Importantly, we also observed an increase in the percentage of T_CM_ cells when the CD4^+^ T cells were transduced with Vpr-expressing lentivirus but not a control lentivirus (Fig 4B, 5 donors combined in Fig S7B). We did not observe this effect in uninfected cells (Fig S7C,D). These data demonstrate that Vpr expression in HIV-1-infected CD4^+^ T cells alters the composition of the overall population by increasing the percentage of cells with a T_CM_ phenotype.

We next examined whether the increase in T_CM_ cells within the culture was affected by any of the Vpr mutations that we had previously examined. Interestingly, none of the four Vpr mutations (F72A/R73A, Q65R, Y50A, and R90K) significantly abrogated the ability of Vpr to increase the proportion of T_CM_ cells in the culture (Fig 4C). These data indicate that the effect of Vpr on the memory phenotype of CD4^+^ T cells is independent of the domains affected by these mutations and is mediated by a distinct mechanism.

### T_CM_ cells are resistant to Vpr-induced apoptosis in HIV-1-infected cells

We hypothesized that the increased fraction of CD4^+^ T cells with a T_CM_ phenotype following Vpr expression could potentially result from one of two mechanisms: 1) Vpr-induced epigenetic and/or transcriptional reprogramming of CD4^+^ T cells to a T_CM_ phenotype or 2) Vpr-mediated killing of non-T_CM_ subsets. To interrogate these possibilities, we examined the level of apoptosis within each T-cell subset (T_N/SCM_, T_CM_, T_EM_ and T_EFF_) after infection with HIV-GFP-Vpr. To identify cells in different stage of apoptosis we stained infected cells at 14 days post-infection with Annexin V and ZV. Cells in early apoptosis were identified as Annexin V^+^/ZV^-^, while cells in late apoptosis/dead were identified as Annexin V^+^/ZV^+^. At 14 days post infection, we observed, as expected, an overall increase in cells that were in both late and early apoptosis for cells infected with HIV-GFP-Vpr but not with HIV-GFP. Interestingly, when we divided the data based on T-cell phenotype, we observed different levels of Annexin V^+^ (apoptotic/dead) cells across subtypes. In particular, cells with a T_CM_ phenotype had lower levels of total and late apoptosis/dead cells than other subtypes (Fig 4D). Interestingly, the level of early apoptotic cells was higher in T_CM_ cells, suggesting that a non-apoptotic pathway may have been responsible for the increased cell death or that most of the apoptosis induced had already occurred by the time we measured cell death/apoptosis at days seven and 14. Importantly, the effects were only observed in productively infected (GFP^+^) cells and were not observed in GFP^-^ cells within the same culture (Fig S4B), demonstrating a cell-intrinsic mechanism for this killing. These data suggest that the ability of Vpr to induce apoptosis/cell death is attenuated in T_CM_ cells compared with other major CD4^+^ T-cell subsets.

To further investigate the effect of Vpr on apoptosis-regulating pathways in infected cells, we also re-examined our bulk RNAseq data from CD4^+^ T cells infected with a Vpr-expressing HIV strain for relative expression of a previously defined array of 44 cell death genes involved in apoptosis, necroptosis, and pyroptosis (53). We hypothesized that, by inducing apoptosis, Vpr expression could potentially select for cells with different levels of expression of these apoptosis-regulating genes. When we compared the transcriptional profiles of cells infected with HIV-GFP-Vpr versus cells infected with HIV-GFP, we observed eight cell death-regulating genes that were differentially expressed: BCL2L11, BIK, BAX, BIRC2, BIRC3, ZBP1, GSDME, and PARP1. We observed upregulation of the necroptosis gene ZBP1, the pro-apoptosis gene BCL2L11, and the pro-pyroptosis gene GSDME in Vpr expressing cells, and we observed downregulation of the pro-apoptotic genes BIK and BAX and upregulation of the caspase antagonists BIRC2 and BIRC3 (Fig S8) in Vpr expressing cells. We speculate that increased expression of BIRC3 and BIRC3 results from selection for cells that express higher levels of these genes. Overall, these data support our hypothesis that the observed enrichment in T_CM_ cells after infection with a Vpr-expressing virus may be due to their ability to resist Vpr-induced apoptosis. Thus, Vpr expression during latency establishment may help to enrich the initial pool of infected cells that enter the reservoir within the T_CM_ compartment.

## Discussion

Recently, it has been appreciated that the a sizable fraction of the long-lived latent reservoir is established or stabilized around the time of ART initiation (19,20,54) Understanding how host and viral factors regulate HIV-1 expression as a provirus enters a transcriptionally silent state, as the infected cell enters the latent reservoir, will lead to identification of candidate therapeutic targets for a latency prevention approach. Whereas factors that maintain proviral quiescence and latency have been studied extensively, the mechanisms involved in latency establishment are less well understood and may differ in some respects from those that maintain latency. We hypothesize that these differences exist both because the host epigenetic and transcriptomic landscapes differ between the two states and because, unlike latent provirus, proviruses undergoing transcriptional downregulation still initially produce viral proteins that can interact with host factors.

We previously identified HDACs as a critical early factor for viral entry into latency, both in terms of transcriptional silencing and for differentiation into T-cell subsets typically enriched for latent virus (21). This prior work focused on host cell factors that regulate HIV-1 silencing in CD4^+^ T cells with the goal of identifying potential cellular targets for interfering with or preventing latency from being established. However, because these experiments relied primarily on a reporter strain of HIV-1 with mutations in the regions encoding the HIV-1 structural and accessory proteins, the role of most HIV-1 genes in transcriptional silencing of HIV-1 in CD4^+^ T cells remained unclear. It is possible that if a specific viral protein that plays an important role in latency establishment can be identified, this protein could also be targeted along with host cell pathways to disrupt latency.

In this study, we identify Vpr as a key viral protein that plays a role in regulating the establishment of a pool of latently infected CD4^+^ T cells and that shapes key characteristics of these cells. Using single gene revertant strains of HIV-1 for Gag/Pol, Vpr, Vpu, Vif, and Nef in a primary CD4^+^ T-cell model of HIV-1 latency establishment, we showed that, while Vpr expression alone does not have a strong effect on viral transcription, Vpr expression in combination with exposure to the class I HDACi vorinostat leads to a dramatic, sustained increase in viral gene expression. Furthermore, we show that this activity depends on a p300-binding domain of Vpr as well as the HAT activity of p300. Additionally, we find that combined Vpr expression and vorinostat exposure induces a potent and unique transcriptional signature in infected cells, further demonstrating the interaction of these pathways. Overall, our results demonstrate that, as HIV-1 enters latency, Vpr has strong pro-transcriptional activity that requires HAT activity of p300 but is blocked by HDACs (Fig 6). Establishment of HIV latency thus results, in significant part, from an HDAC-mediated restriction to Vpr activity in CD4 T cells.

**Fig 6.**
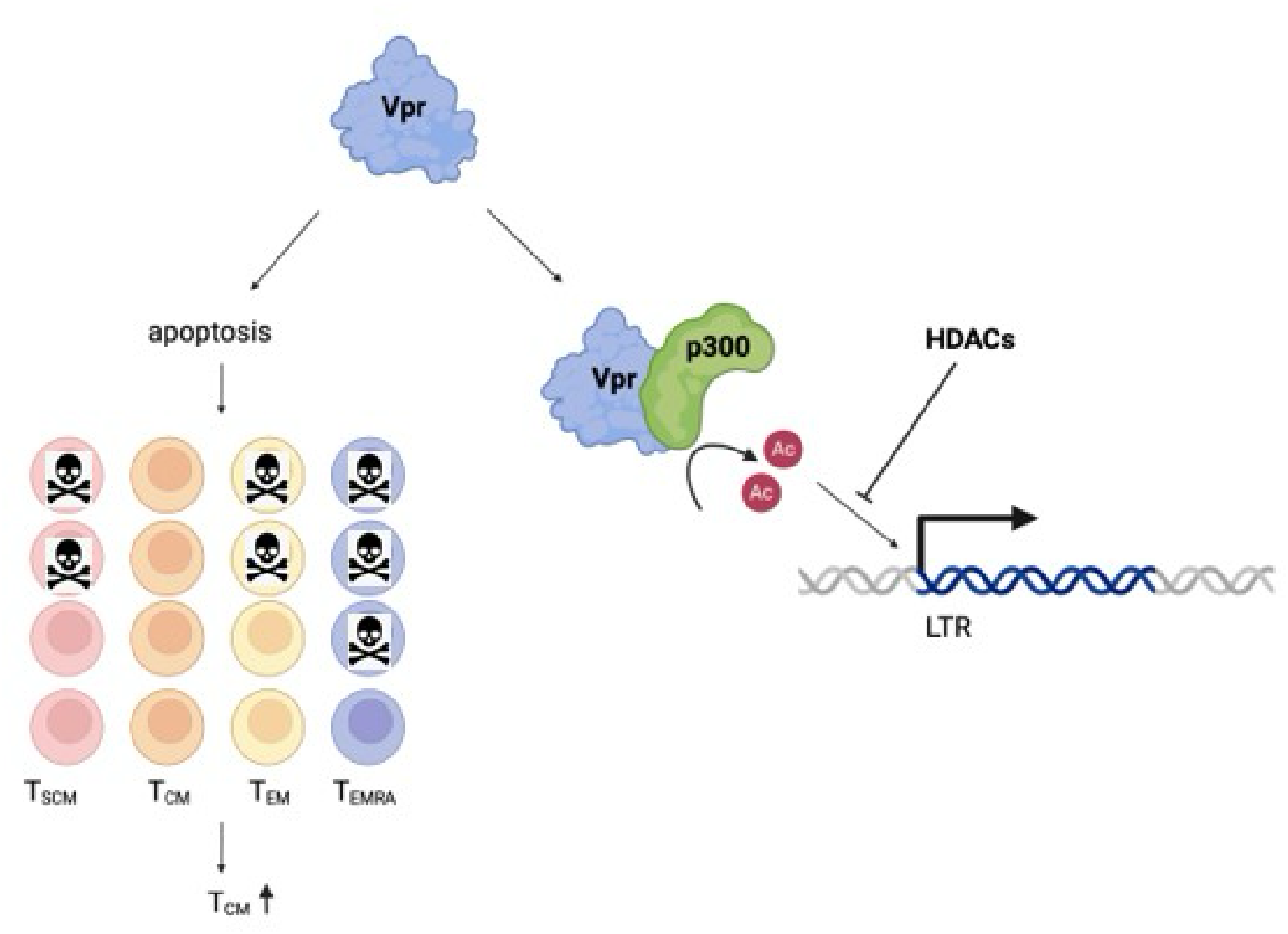
Model of role of Vpr in HIV-1 transcription and T-cell memory subset composition. Left: Vpr in memory CD4+ T-cell subset composition. Vpr expression leads to enrichment for TCM-like cells (CD45RA^-^CCR7^+^), likely because of selective cell death by other memory compartments. Right: Vpr in HIV-1 transcription. Vpr transactivates the HIV-1 promoter (long terminal repeat; LTR) in a manner dependent on an intact p300-binding domain as well as an intact p300 acetyltransferrase domain. Histone acetyltransferrase activity blocks this pro-transcriptional activity of Vpr. Created with Biorender.com

Previous studies have indicated a pro-transcriptional activity for Vpr (28,29,31,46,55,56). However, the mechanism involved is unclear and has been attributed to multiple functions of Vpr, most commonly cell cycle arrest (29,31). Studies have also reported a link between Vpr-induced cell cycle arrest and DCAF1-binding and pro-apoptotic activity, making it difficult to distinguish each function’s individual contribution to viral promoter transactivation. Vpr has also been shown to transactivate HIV-1 gene expression via p300, both directly and indirectly (28,46,57), although this function is less well-studied. Many of these previous Vpr studies were performed in immortalized cell lines. Although cell lines have been an invaluable tool for studying many aspects of HIV-1 virology and pathogenesis, they have been modified to evade cellular senescence, making them unlikely to be the most accurate model of an HIV-1-infected cell returning to rest. Further, previous studies have often relied solely on lentiviral expression vectors with strong promoters driving Vpr and/or expression during HIV-latency. This can lead to nonbiological levels of Vpr expression in an epigenetic or transcriptional landscape in which viral protein would not be produced in sizable amounts. To address these challenges, we infected primary CD4^+^ T cells with virus in which Vpr expression was driven by the viral promoter or observed the action of exogenous Vpr on the viral promoter.

The p300 HAT domain has previously been shown to activate HIV-1 gene expression via several possible mechanisms, including histone acetylation-mediated chromatin remodeling and acetylation of the viral protein Tat and host transcription factors such as NF-κB (58). Tat acetylation is required for Tat-mediated transactivation of the HIV-1 promoter. Acetylation of NF-κB subunits enables NF-κB to bind with more stability to NF-κB-binding sites and to the basal transcription machinery, promoting HIV-1 gene expression (58,59). However, further study will be required to better understand the role of Vpr, HDACs, and p300 HAT activity in regulating HIV-1 gene expression, particularly as the virus enters transcriptional latency. These studies will be challenging because the toxicity of Vpr limits the number of cells available for analyses. Understanding how Vpr regulates HIV-1 transcription in the context of host epigenetic and transcription factors will aid in the development of agents that can prevent latency formation or more potently reverse latency. Identifying the individual HDAC that counteracts Vpr’s pro-transcriptional activity in T cells will also help inform therapeutic approaches to targeting the reservoir.

Following cis or trans Vpr expression, we also observed an increase in cells with a T_CM_-like phenotype (CCR7^+^CD45RA^-^; Fig 4A,B, Fig S7A,B), a long-lived cell type in which the a significant fraction of the latent reservoir is thought to reside (24,60,61). Our findings suggest that Vpr, which is packaged within the virion and therefore present from the early stages of infection (62,63), could play an important role in shaping the initial location of latent reservoir. Although our data are consistent with a “selection” model for how Vpr increases the relative frequency of T_CM_ cells in the latent pool, it is also possible that Vpr reprogramming of CD4^+^ T cells into T_CM_-like cells occurs. Previous work has shown that the majority of the early transcriptional (64) as well as the proteomic changes (65) that occur following HIV-1 infection are driven by Vpr. In addition, in resting CD4^+^ T cells infected by cell-to-cell transmission, Vpr reprogrammed cells to a tissue resident memory phenotype (66). However, in our bulk RNASeq dataset, none of the pathways enriched by GSEA using the Hallmark gene set would clearly lead to memory cell reprogramming (Fig S7F). Our data instead support a model in which T_CM_ cells are more resistant to Vpr-induced apoptosis/cell death than other memory cell subsets, leading to selective enrichment of infected T_CM_ cells in the overall pool of infected cells (Fig 6). Indeed, previous work has shown that T_CM_ cells are less susceptible to cell death following HIV reactivation (67,68). Interestingly, we did not observe a change in the enrichment for T_CM_-like cells following infection with any of the four Vpr mutants generated, including the mutant that lacks apoptosis-inducing ability (Fig 4C), indicating that the preferential survival of T_CM_ cells may be a complex phenomenon. We also speculate that, when viral gene expression from latent proviruses is reignited, either sporadically or after LRA-mediated reactivation, infected T_CM_ cells are preferentially protected from Vpr-mediated cell death, allowing these cells to persist despite effective latency reversal. Indeed, even LRAs that have shown potent latency-reversing activity in vivo have had no apparent effect on the size of the reservoir (15,69–71). Thus, the resistance of T_CM_ cells to Vpr-mediated killing could represent a key mechanism of viral persistence.

The mechanism by which T_CM_ cells resist Vpr-induced apoptosis/cell death is still unclear. In our bulk RNAseq data, we observed an upregulation of two caspase antagonists (BIRC2, BIRC3) as well as downregulation of two pro-apoptotic proteins (BIK, BAX) in cells that had survived infection with a Vpr-expressing virus for two weeks, while a different apoptosis gene (BCL2L11), a pro-necroptosis pathway gene (ZBP1), and a pro-pyroptosis gene (GSDME) were upregulated (Fig S8). This differential gene expression suggests that Vpr induces necroptosis, pyroptosis, and apoptosis in our model but that T_CM_ cells may express higher levels of caspase antagonists, which protects them from apoptosis. Interestingly, a previous study implicated the BIRC2 and BIRC3 protein products cIAP1 and cIAP3, respectively, in protecting macrophages from Vpr-induced cell death during HIV-1 infection (72).

It is important to note that the observed BIRC2/3 upregulation in our study is from bulk RNA sequencing. Additional study on sorted memory cell populations or single-cell RNA sequencing may therefore be informative. One new approach to targeting persistent, long-lived memory cell populations has been to sensitize cells to apoptosis by using the BCL-2 antagonist venetoclax (68). However, in cells from PWH treated with venetoclax ex vivo, T_CM_ cells were largely spared, suggesting that T_CM_ cells possess unique mechanisms of resisting apoptosis. Understanding how Vpr affects survival and how the latent reservoir forms has important implications both for potentially preventing the effector-to-memory transition that occurs during ART initiation and for targeting latently infected memory cells for cell death (67,73).

## Limitations of the study

We acknowledge several limitations of the study. First, the use of single gene revertants enables the effects of individual proteins on gene expression and CD4^+^ T-cell memory subset composition to be separated but did not allow us to study the combined effect of HIV-1 Gag/Pol and accessory proteins. It is possible that wild type Vpr is expressed at higher levels than the Vpr revertant virus in our model. We also observed significant overlap in the phenotypes of our mutant Vpr viruses, for example, cell death/apoptosis was not only decreased in the R90K mutant but also to some extent in the other mutants as well. It is possible that future work would benefit from including additional Vpr mutants. However, studies have shown that the functions of Vpr may be interrelated (36,49,74), suggesting that it may be challenging to generate single-function Vpr mutant viruses. It is also important to note that the p300 inhibitors used were not specific to Vpr-p300 interaction and some of the observed effect on transcription may have been Vpr-independent. The lack of effect on GFP expression in either HIV-GFP- or HIV-GFP-Vpr (F72A/R73A)-infected cells indicates though that the abrogation of HIV expression in HIV-GFP-Vpr-infected cells is Vpr-dependent. Finally, for all experiments, cells were activated ex vivo with anti-CD3/CD28 antibodies and maintained in culture under IL-2/IL-7 conditions, which is likelier a stronger stimulation than most T cells experience in vivo and could alter memory formation. In addition, only two surface markers (CCR7 and CD45RA) were used to define memory cell subset. Future work will benefit from including additional surface markers and/or performing functional assays.

## Methods and Materials

### Plasmids and viruses

We modified a previously published HIV reporter virus NL4-3-D6-dreGFP (25), herein referred to as HIV-GFP. This strain of HIV contains inactivating mutations in regions encoding all viral proteins except Tat and Rev. In the parental plasmid, the *nef* gene is truncated and point mutations have been introduced to generate premature stop codons in *gag*, *vif*, *vpr*, and *vpu*. A portion of the envelope has also been replaced with destabilized eGFP (dreGFP), and we previously inserted an IRES-Thy1.2 cassette immediately downstream of dreGFP (21) (Fig 1B). Because Thy1.2 has a significantly longer half-life than dreGFP, these two markers can be used to differentiate between productively infected cells (Thy1.2^+^GFP^+^) and infected cells that have recently downregulated HIV expression (Thy1.2^+^GFP^-^). We used site-directed mutagenesis to individually revert the stop codons in *gag/pol, vif, vpr, vpu.* To restore *nef* expression, we digested the HIV-GFP plasmid with BamH1 and Xho1 and inserted a gblock (IDT) with the missing *nef* sequence to restore the complete open reading frame. Lentiviral expression vectors (Addgene 21373) were generated by replacing the GFP reporter gene with mCherry and inserting Vpr coding sequence into the multiple cloning site. Mutated Vpr constructs (Y50A, R90K, Q65R, F72A/R73A) in both the HIV-GFP and lentiviral expression vector backgrounds were generated using NEB assembly. Full plasmid sequences of Vpr mutant plasmids were confirmed by sequencing.

### Virus production

Virus stocks were generated by transfecting HEK293T/17 cells (ATCC) with packaging plasmids psPAX2 and MD2-VSVG in combination with the relevant viral backbone plasmid using Mirus LT1 transfection reagent at a 3:1 volume to mass ratio. DMEM (Gibco; supplemented with 10% fetal calf serum (FCS) and 1% Penicillin Streptomycin) was replaced with RPMI R10 (10% FCS; 1% Penicillin Streptomycin; 10mM HEPES; 2mM L-Glutamine; 1mM Sodium Pyruvate) 24h post-transfection, and virus-containing supernatant was collected 48h post-transfection. Supernatant was then centrifuged at low speed to remove cellular debris, filtered through a 0.45μm low protein-binding filter, and frozen at -80°C before use.

### Primary CD4^+^ T-cell infection model and cell culture

CD4^+^ T cells were obtained from leuko-reduction 769 blood products (STEMCELL Technologies) by gradient centrifugation followed by magnetic isolation and cyropreserved in freezing medium (90% FCS, 10% DMSO) as previously described (21). Cells were thawed and activated using anti-CD3/anti-CD28 Dynabeads (Gibco) in RPMI R10 supplemented with 100U/mL IL-2 and 1ng/mL IL-7 (Peprotech). After 48h, the activating beads were removed, and the cells were infected by resuspending in media containing virus stocks with 4μg/mL polybrene (Hexadimethrine Bromide) and then spinoculated for 2h at 600g. Viral supernatant was then replaced with RPMI at a concentration of 10^6^ cells/mL. Active infection (%GFP^+^ cells within Thy1.2+ gate) was quantified by flow cytometry 48-72h after infection. Media containing appropriate compound (500nM vorinostat, DMSO vehicle control, or p300 inhibitor as detailed below) was replaced every 2-3 days, and cell density was increased to 2x10^6^ cells/mL after 7 days. Productive infection (%GFP^+^ within Thy1.2^+^), viral gene expression (median fluorescence intensity; MFI of productively infected cells), memory phenotype (by CCR7 and CD45RA expression), apoptosis (Annexin V and Zombie-NIR), and cell cycle 4′,6-diamidino-2-phenylindole (DAPI) were measured by flow cytometry every 7 days (Fig 1A).

### Flow cytometry

Cells were washed with phosphate buffered saline (PBS) and stained with Live-Dead viability dye (Zombie Violet or Zombie-NIR, BioLegend) and an anti-CD90.2 antibody (Thy1.2, BioLegend) in PBS. For experiments assessing cell surface expression of memory markers, cells were then washed with FACS buffer (PBS with 2% 776 FCS, 1mM EDTA) and stained with antibodies to CD45RA and CCR7 (BioLegend). For experiments assessing apoptosis, cells were stained with Annexin V in Annexin V binding buffer (BioLegend) and analyzed by flow cytometry without fixation. For all other experiments, cells were fixed in 4% paraformaldehyde (PFA) prior to acquisition on a Fortessa flow cytometer (Becton Dickson). Data were analyzed using FlowJo (version 10.1). To quantify cells in G2/M phase, cells were stained post-fixation with DAPI (BioLegend) diluted to 3μM in 1x FOXP3 perm buffer (BioLegend). For GFP quantification, at least 10,000 live cells were collected for analysis. To assess memory markers, at least 50,000 live cells were recorded. Compensation controls were prepared using single-color stained cells. To compensate for the viability stain and CD90.2 antibody, ArC™ amine reactive compensation beads and UltraComp eBeads Plus compensation beads were used, respectively (Invitrogen). Positive events were gated using fluorescence minus one and biological (uninfected cells) controls.

### Gag transcript quantification

Fourteen days post-infection (14dpi) with either HIV-GFP or the HIV-GFP-Vpr revertant, GFP^+^ CD4^+^ T cells from two donors, treated with either DMSO or vorinostat, were sorted into RLT^+^ lysis buffer (Qiagen). RNA was isolated using RNeasy Plus Mini Kit (Qiagen 74134), and Gag transcript was quantified by RT-qPCR on a QuantStudio 3 Real-Time PCR system (Thermo Fisher Scientific) using *β*-actin as a reference gene. A commercially purchased Taqman primer/probe set was used for *β*-actin (Thermo Fisher Scientific Hs01060665_g1). Double delta CT analysis was used to calculate fold-change in *Gag* transcript levels.

### p300 inhibitors

Two different p300 inhibitors were used: bromodomain inhibitor (CCS1477, Selleckchem) and histone acetyltransferase inhibitor (A-485, Selleckchem). Prior to experiments in infected cells, dose curves were performed to assess toxicity and cell viability. Based both on viability and previously reported EC_50_, 0.122μM CCS1477 and 0.56nM A-485 were selected (50,51).

### Western blotting

To generate whole cell protein lysates, HEK293T/17 cells were harvested and lysed with RIPA buffer (Thermo Fisher) supplemented with 1x protease inhibitors (Roche) and 1% Pierce Universal Nuclease (Thermo Fisher). Protein quantification was performed using DC Protein Assay (Bio-Rad). Proteins were resolved by Tris-Glycine or Tris-Acetate SDS-PAGE depending on protein size and transferred to nitrocellulose membrane. Membranes were blocked in 5% milk in 1x Tris-Buffered Saline (TBS) and probed at 4°C overnight with antibodies (Vpr, Proteintech 51143-1-AP; Hsp90, Cell Signaling Technology 4877) diluted in TBS containing 5% BSA or milk. Membranes were washed with TBS containing 0.1% Tween20 (TBST), probed with secondary antibody (1:10,000) conjugated to horseradish peroxidase for 1h at room temperature, washed again with TBST, and developed using enhanced chemiluminescence (Thermo Fisher).

### RNA sequencing

RNA isolation was performed with RNeasy Plus Mini Kit (Qiagen 74134), and cDNA synthesis and library prep were performed using SMART-Seq mRNA LP (Takara 634768). Libraries were quality controlled and quantified by Qubit dsDNA assay (Thermo Fisher) and capillary electrophoresis (Agilent Tapestation) prior to pooling and 2x50 paired-end sequencing on an Illumina NextSeq 2000 system. Quality reports were generated by fastQC v 0.11.9, and reads were aligned to the GRCh38 human genome as well as a previously generated custom HIV reference genome containing the HIV-GFP sequence using STAR v2.7.10b. Differential gene expression was assessed in R using DESeq2 v1.40.2 (75).

### Statistics

Statistical analyses were performed in GraphPad Prism and R. To analyze differences in GFP MFI and within memory subsets between HIV-GFP-Vpr and HIV-GFP, Wilcoxon matched-pairs signed-rank test was employed for studies that were sufficiently powered (n > 6). Benjamini and Hochberg correction with a false-discovery rate of 0.05 was used.

## Acknowledgements

We would like to thank Kimberly Enders at the Center for AIDs Research at UNC for her guidance with statistical analysis. We would also like to thank Anne-Marie Turner and Joshua Fox at the Institute for Global Health and Infectious Diseases for performing Oxford Nanopore sequencing on our Vpr mutant plasmids. Finally, we would like to acknowledge the UNC Flow Cytometry Core Facility (RRID:SCR_019170).

## Supporting information captions

**Fig S1. HIV-GFP revertant viruses.** A) Western blot for Vif, Vpr, and p24 of 293T cells transfected with HIV-GFP revertant plasmids. B) CD4 expression in CD4+ T cells infected with HIV-GFP revertant viruses. C) Quantification of GFP+ CD4+ T cells within Thy1.2+ population for each virus. D) Representative flow plots for infection with each revertant virus. GFP+ cells were gated within Thy1.2+ cells.

**Fig S2.** Infection of CD4+ T cells with single gene revertant viruses. Combined data for multiple donors. Each donor is represented by a different symbol. A-B) GFP median fluorescence intensity (MFI) from A) cis Vpr expression (n = 9 for weeks 1,2 and n = 6 for week 3) and D) trans Vpr expression (n = 6 for weeks 1, 2 and n = 2 for week 3). C-D) Gag unspliced RNA and GFP MFI within GFP+ cells for C) HIV-GFP-Vpr vs. HIV-GFP (n = 2 donors) and D) cells co-infected with HIV-GFP and Vpr-mCherry or mCherry (n = 3 donors).

**Fig S3.** Vpr localization within cell compartments does not change following vorinostat treatment. Western blot with indicated antibody of cell fractions from cell treated with DMSO or vorinostat. VOR = vorinostat. Sol nuc = soluble nuclear fraction.

**Fig S4.** Induction of cell cycle arrest and apoptosis/cell death by Vpr. A) Left: example histogram of DAPI staining for cell cycle. Right: % cells in G2/M in each condition 7 days post-infection. B) Representative flow plots of cells stained with Annexin V and Zombie Violet to measure apoptosis and cell death 14 days post-infection.

**Fig S5.** Induction of cell cycle arrest and apoptosis/cell death by Vpr functional mutant viruses. Vpr expression in 293T cells transfected with mutant A) HIV-GFP-Vpr and B) Vpr-mCherry plasmids.

**Fig S6.** Combined results from p300 inhibitor experiments (n = 3). CD4+ T cells infected with HIV-GFP, HIV-GFP-Vpr, or HIV-GFP-Vpr (F72/F73A) virus and treated with DMSO, VOR, or A) p300 bromodomain inhibitor (CCS1477) alone or in combination with VOR or B) p300 histone acetyltransferrase inhibitor (A485) alone or in combination with VOR. Each symbol represents a different donor.

**Fig S7.** Memory subset composition. Combined data from multiple donors for %GFP+ (productively infected) cells in each subset as defined by CCR7 and CD45RA in cells infected with A) HIV-GFP-Vpr or HIV-GFP (n = 6) or B) Vpr-mCherry or mCherry lentivirus (n = 5). Each symbol represents a different donor. C-D) Representative from same experiments of %GFP-(uninfected) cells. E) Heat map of differentially expressed genes (DEGs) in GFP+ CD4+ T cells infected with HIV-GFP-Vpr or HIV-GFP (n = 3). F) gene-set enrichment analysis for all DEGs.

**Fig S8.** DEGs in HIV-GFP-Vpr- vs. HIV-GFP-infected cells within a previously published subset of cell death pathway-related genes (53).

## References

1. Kotey M, Alhassan Y, Adomako J, Nunoo-Mensah G, Kapadia F, Sarfo B. Chronic comorbidities in persons living with HIV within three years of exposure to antiretroviral therapy at Pantang Antiretroviral Center in Ghana: a retrospective study. Pan Afr Med J. 2022 Aug 19;42:294.

2. Pourcher V, Gourmelen J, Bureau I, Bouee S. Comorbidities in people living with HIV: An epidemiologic and economic analysis using a claims database in France. PLoS One. 2020 Dec 17;15(12):e0243529.

3. Armstrong-Mensah EA, Tetteh AK, Ofori E, Ekhosuehi O. Voluntary Counseling and Testing, Antiretroviral Therapy Access, and HIV-Related Stigma: Global Progress and Challenges. Int J Environ Res Public Health. 2022 May 28;19(11):6597.

4. Oturu K, O’Brien O, Ozo-Eson PI. Barriers and enabling structural forces affecting access to antiretroviral therapy in Nigeria. BMC Public Health. 2024 Jan 6;24(1):105.

5. Barnabas RV, Szpiro AA, Ntinga X, Mugambi ML, Rooyen H van, Bruce A, et al. Fee for home delivery and monitoring of antiretroviral therapy for HIV infection compared with standard clinic-based services in South Africa: a randomised controlled trial. The Lancet HIV. 2022 Dec 1;9(12):e848–56.

6. Siliciano JD, Kajdas J, Finzi D, Quinn TC, Chadwick K, Margolick JB, et al. Long-term follow-up studies confirm the stability of the latent reservoir for HIV-1 in resting CD4+ T cells. Nat Med. 2003 Jun;9(6):727–8.

7. Crooks AM, Bateson R, Cope AB, Dahl NP, Griggs MK, Kuruc JD, et al. Precise Quantitation of the Latent HIV-1 Reservoir: Implications for Eradication Strategies. The Journal of Infectious Diseases. 2015 Nov 1;212(9):1361–5.

8. McMyn NF, Varriale J, Fray EJ, Zitzmann C, MacLeod H, Lai J, et al. The latent reservoir of inducible, infectious HIV-1 does not decrease despite decades of antiretroviral therapy. J Clin Invest. 133(17):e171554.

9. Coffin JM, Wells DW, Zerbato JM, Kuruc JD, Guo S, Luke BT, et al. Clones of infected cells arise early in HIV-infected individuals. JCI Insight. 2019 Jun 20;4(12):e128432, 128432.

10. Chun TW, Stuyver L, Mizell SB, Ehler LA, Mican JAM, Baseler M, et al. Presence of an inducible HIV-1 latent reservoir during highly active antiretroviral therapy. Proceedings of the National Academy of Sciences. 1997 Nov 25;94(24):13193–7.

11. Wong JK, Hezareh M, Günthard HF, Havlir DV, Ignacio CC, Spina CA, et al. Recovery of Replication-Competent HIV Despite Prolonged Suppression of Plasma Viremia. Science. 1997 Nov 14;278(5341):1291–5.

12. Finzi D, Hermankova M, Pierson T, Carruth LM, Buck C, Chaisson RE, et al. Identification of a Reservoir for HIV-1 in Patients on Highly Active Antiretroviral Therapy. Science. 1997 Nov 14;278(5341):1295–300.

13. Uldrick TS, Adams SV, Fromentin R, Roche M, Fling SP, Gonçalves PH, et al. Pembrolizumab induces HIV latency reversal in people living with HIV and cancer on antiretroviral therapy. Science Translational Medicine. 2022 Jan 26;14(629):eabl3836.

14. Søgaard OS, Graversen ME, Leth S, Olesen R, Brinkmann CR, Nissen SK, et al. The Depsipeptide Romidepsin Reverses HIV-1 Latency In Vivo. PLOS Pathogens. 2015 Sep 17;11(9):e1005142.

15. Archin NM, Liberty AL, Kashuba AD, Choudhary SK, Kuruc JD, Crooks AM, et al. Administration of vorinostat disrupts HIV-1 latency in patients on antiretroviral therapy. Nature. 2012 Jul;487(7408):482–5.

16. Archin NM, Cheema M, Parker D, Wiegand A, Bosch RJ, Coffin JM, et al. Antiretroviral intensification and valproic acid lack sustained effect on residual HIV-1 viremia or resting CD4+ cell infection. PLoS One. 2010 Feb 23;5(2):e9390.

17. Archin NM, Kirchherr JL, Sung JAM, Clutton G, Sholtis K, Xu Y, et al. Interval dosing with the HDAC inhibitor vorinostat effectively reverses HIV latency. J Clin Invest. 127(8):3126– 35.

18. Nixon CC, Mavigner M, Sampey GC, Brooks AD, Spagnuolo RA, Irlbeck DM, et al. Systemic HIV and SIV latency reversal via non-canonical NF-κB signalling in vivo. Nature. 2020 Feb;578(7793):160–5.

19. Abrahams MR, Joseph SB, Garrett N, Tyers L, Moeser M, Archin N, et al. The replication-competent HIV-1 latent reservoir is primarily established near the time of therapy initiation. Sci Transl Med. 2019 Oct 9;11(513):eaaw5589.

20. Brodin J, Zanini F, Thebo L, Lanz C, Bratt G, Neher RA, et al. Establishment and stability of the latent HIV-1 DNA reservoir. Chakraborty AK, editor. eLife. 2016 Nov 15;5:e18889.

21. Peterson JJ, Lewis CA, Burgos SD, Manickam A, Xu Y, Rowley AA, et al. A histone deacetylase network regulates epigenetic reprogramming and viral silencing in HIV-infected cells. Cell Chem Biol. 2023 Dec 21;30(12):1617–1633.e9.

22. Sengupta S, Siliciano RF. Targeting the latent reservoir for HIV-1. Immunity. 2018 May 15;48(5):872–95.

23. Peterson JJ, Lewis CA, Burgos SD, Manickam A, Xu Y, Rowley AA, et al. A histone deacetylase network regulates epigenetic reprogramming and viral silencing in HIV infected cells [Internet]. bioRxiv; 2022 [cited 2023 May 30]. p. 2022.05.09.491199. Available from: https://www.biorxiv.org/content/10.1101/2022.05.09.491199v1

24. Soriano-Sarabia N, Bateson RE, Dahl NP, Crooks AM, Kuruc JD, Margolis DM, et al. Quantitation of replication-competent HIV-1 in populations of resting CD4+ T cells. J Virol. 2014 Dec;88(24):14070–7.

25. Yang HC, Xing S, Shan L, O’Connell K, Dinoso J, Shen A, et al. Small-molecule screening using a human primary cell model of HIV latency identifies compounds that reverse latency without cellular activation. J Clin Invest. 2009 Nov 2;119(11):3473–86.

26. Jefferys SR, Burgos SD, Peterson JJ, Selitsky SR, Turner AMW, James LI, et al. Epigenomic characterization of latent HIV infection identifies latency regulating transcription factors. PLoS Pathog. 2021 Feb;17(2):e1009346.

27. Romani B, Kamali Jamil R, Hamidi-Fard M, Rahimi P, Momen SB, Aghasadeghi MR, et al. HIV-1 Vpr reactivates latent HIV-1 provirus by inducing depletion of class I HDACs on chromatin. Sci Rep. 2016 Aug 23;6(1):31924.

28. Kino T, Gragerov A, Slobodskaya O, Tsopanomichalou M, Chrousos GP, Pavlakis GN. Human Immunodeficiency Virus Type 1 (HIV-1) Accessory Protein Vpr Induces Transcription of the HIV-1 and Glucocorticoid-Responsive Promoters by Binding Directly to p300/CBP Coactivators. J Virol. 2002 Oct;76(19):9724–34.

29. Goh WC, Rogel ME, Kinsey CM, Michael SF, Fultz PN, Nowak MA, et al. HIV-1 Vpr increases viral expression by manipulation of the cell cycle: A mechanism for selection of Vpr in vivo. Nat Med. 1998 Jan;4(1):65–71.

30. Yao XJ, Mouland AJ, Subbramanian RA, Forget J, Rougeau N, Bergeron D, et al. Vpr Stimulates Viral Expression and Induces Cell Killing in Human Immunodeficiency Virus Type 1-Infected Dividing Jurkat T Cells. Journal of Virology. 1998 Jun;72(6):4686–93.

31. Gummuluru S, Emerman M. Cell Cycle- and Vpr-Mediated Regulation of Human Immunodeficiency Virus Type 1 Expression in Primary and Transformed T-Cell Lines. J Virol. 1999 Jul;73(7):5422–30.

32. Zhang F, Bieniasz PD. HIV-1 Vpr induces cell cycle arrest and enhances viral gene expression by depleting CCDC137. eLife. 9:e55806.

33. Depienne C, Roques P, Créminon C, Fritsch L, Casseron R, Dormont D, et al. Cellular Distribution and Karyophilic Properties of Matrix, Integrase, and Vpr Proteins from the Human and Simian Immunodeficiency Viruses. Experimental Cell Research. 2000 Nov 1;260(2):387–95.

34. Muthumani K, Choo AY, Zong WX, Madesh M, Hwang DS, Premkumar A, et al. The HIV-1 Vpr and glucocorticoid receptor complex is a gain-of-function interaction that prevents the nuclear localization of PARP-1. Nat Cell Biol. 2006 Feb;8(2):170–9.

35. de Noronha CMC, Sherman MP, Lin HW, Cavrois MV, Moir RD, Goldman RD, et al. Dynamic Disruptions in Nuclear Envelope Architecture and Integrity Induced by HIV-1 Vpr. Science. 2001 Nov 2;294(5544):1105–8.

36. Stewart SA, Poon B, Jowett JB, Chen IS. Human immunodeficiency virus type 1 Vpr induces apoptosis following cell cycle arrest. J Virol. 1997 Jul;71(7):5579–92.

37. Jowett JB, Planelles V, Poon B, Shah NP, Chen ML, Chen IS. The human immunodeficiency virus type 1 vpr gene arrests infected T cells in the G2 + M phase of the cell cycle. J Virol. 1995 Oct;69(10):6304–13.

38. Bartz SR, Rogel ME, Emerman M. Human immunodeficiency virus type 1 cell cycle control: Vpr is cytostatic and mediates G2 accumulation by a mechanism which differs from DNA damage checkpoint control. J Virol. 1996 Apr;70(4):2324–31.

39. Bernhart E, Stuendl N, Kaltenegger H, Windpassinger C, Donohue N, Leithner A, et al. Histone deacetylase inhibitors vorinostat and panobinostat induce G1 cell cycle arrest and apoptosis in multidrug resistant sarcoma cell lines. Oncotarget. 2017 Sep 29;8(44):77254– 67.

40. Duvic M, Vu J. Vorinostat in cutaneous T-cell lymphoma. Drugs Today (Barc). 2007 Sep;43(9):585–99.

41. Xu J, Sampath D, Lang FF, Prabhu S, Rao G, Fuller GN, et al. Vorinostat modulates cell cycle regulatory proteins in glioma cells and human glioma slice cultures. J Neurooncol. 2011 Nov;105(2):241–51.

42. Exposure of phosphatidylserine on the surface of apoptotic lymphocytes triggers specific recognition and removal by macrophages. | The Journal of Immunology | American Association of Immunologists [Internet]. [cited 2024 Jul 18]. Available from: https://journals.aai.org/jimmunol/article/148/7/2207/25274/Exposure-of-phosphatidylserine-on-the-surface-of

43. Moon HS, Yang JS. Role of HIV Vpr as a Regulator of Apoptosis and an Effector on Bystander Cells. Molecules and Cells. 2006 Feb 1;21(1):7–20.

44. Stewart SA, Poon B, Song JY, Chen ISY. Human Immunodeficiency Virus Type 1 Vpr Induces Apoptosis through Caspase Activation. J Virol. 2000 Apr;74(7):3105–11.

45. Muthumani K, Hwang DS, Desai BM, Zhang D, Dayes N, Green DR, et al. HIV-1 Vpr Induces Apoptosis through Caspase 9 in T Cells and Peripheral Blood Mononuclear Cells *. Journal of Biological Chemistry. 2002 Oct 4;277(40):37820–31.

46. Felzien LK, Woffendin C, Hottiger MO, Subbramanian RA, Cohen EA, Nabel GJ. HIV transcriptional activation by the accessory protein, VPR, is mediated by the p300 co-activator. Proc Natl Acad Sci U S A. 1998 Apr 28;95(9):5281–6.

47. Maudet C, Bertrand M, Le Rouzic E, Lahouassa H, Ayinde D, Nisole S, et al. Molecular Insight into How HIV-1 Vpr Protein Impairs Cell Growth through Two Genetically Distinct Pathways. J Biol Chem. 2011 Jul 8;286(27):23742–52.

48. Jacquot G, Le Rouzic E, David A, Mazzolini J, Bouchet J, Bouaziz S, et al. Localization of HIV-1 Vpr to the nuclear envelope: impact on Vpr functions and virus replication in macrophages. Retrovirology. 2007 Nov 26;4:84.

49. Le Rouzic E, Belaïdouni N, Estrabaud E, Morel M, Rain JC, Transy C, et al. HIV1 Vpr Arrests the Cell Cycle by Recruiting DCAF1/VprBP, a Receptor of the Cul4-DDB1 Ubiquitin Ligase. Cell Cycle. 2007 Jan 15;6(2):182–8.

50. Welti J, Sharp A, Brooks N, Yuan W, McNair C, Chand SN, et al. Targeting the p300/CBP Axis in Lethal Prostate Cancer. Cancer Discovery. 2021 May 1;11(5):1118–37.

51. Lasko LM, Jakob CG, Edalji RP, Qiu W, Montgomery D, Digiammarino EL, et al. Discovery of a potent catalytic p300/CBP inhibitor that targets lineage-specific tumors. Nature. 2017 Oct 5;550(7674):128–32.

52. Liberzon A, Birger C, Thorvaldsdóttir H, Ghandi M, Mesirov JP, Tamayo P. The Molecular Signatures Database Hallmark Gene Set Collection. Cell Systems. 2015 Dec 23;1(6):417– 25.

53. Rich AL, Lin P, Gamazon ER, Zinkel SS. The broad impact of cell death genes on the human disease phenome. Cell Death Dis. 2024 Apr 8;15(4):1–12.

54. Jones BR, Kinloch NN, Horacsek J, Ganase B, Harris M, Harrigan PR, et al. Phylogenetic approach to recover integration dates of latent HIV sequences within-host. Proceedings of the National Academy of Sciences. 2018 Sep 18;115(38):E8958–67.

55. Cohen EA, Terwilliger EF, Jalinoos Y, Proulx J, Sodroski JG, Haseltine WA. Identification of HIV-1 vpr product and function. J Acquir Immune Defic Syndr (1988). 1990;3(1):11–8.

56. Wang L, Mukherjee S, Jia F, Narayan O, Zhao LJ. Interaction of virion protein Vpr of human immunodeficiency virus type 1 with cellular transcription factor Sp1 and trans-activation of viral long terminal repeat. J Biol Chem. 1995 Oct 27;270(43):25564–9.

57. Kino T, Gragerov A, Kopp JB, Stauber RH, Pavlakis GN, Chrousos GP. The HIV-1 Virion-associated Protein Vpr Is a Coactivator of the Human Glucocorticoid Receptor. Journal of Experimental Medicine. 1999 Jan 4;189(1):51–62.

58. Ott M, Dorr A, Hetzer-Egger C, Kaehlcke K, Schnolzer M, Henklein P, et al. Tat Acetylation: A Regulatory Switch between Early and Late Phases in HIV Transcription Elongation. In: Reversible Protein Acetylation [Internet]. John Wiley & Sons, Ltd; 2004 [cited 2024 Dec 1]. p. 182–96. Available from: https://onlinelibrary.wiley.com/doi/abs/10.1002/0470862637.ch13

59. Mukherjee SP, Behar M, Birnbaum HA, Hoffmann A, Wright PE, Ghosh G. Analysis of the RelA:CBP/p300 interaction reveals its involvement in NF-κB-driven transcription. PLoS Biol. 2013 Sep;11(9):e1001647.

60. Chomont N, El-Far M, Ancuta P, Trautmann L, Procopio FA, Yassine-Diab B, et al. HIV reservoir size and persistence are driven by T cell survival and homeostatic proliferation. Nat Med. 2009 Aug;15(8):893–900.

61. Zerbato JM, Serrao E, Lenzi G, Kim B, Ambrose Z, Watkins SC, et al. Establishment and Reversal of HIV-1 Latency in Naive and Central Memory CD4+ T Cells In Vitro. J Virol. 2016 Aug 26;90(18):8059–73.

62. Hrimech M, Yao XJ, Bachand F, Rougeau N, Cohen ÉA. Human Immunodeficiency Virus Type 1 (HIV-1) Vpr Functions as an Immediate-Early Protein during HIV-1 Infection. J Virol. 1999 May;73(5):4101–9.

63. Human immunodeficiency virus vpr product is a virion-associated regulatory protein [Internet]. [cited 2023 Sep 17]. Available from: https://journals.asm.org/doi/epdf/10.1128/jvi.64.6.3097-3099.1990

64. Bauby H, Ward CC, Hugh-White R, Swanson CM, Schulz R, Goujon C, et al. HIV-1 Vpr Induces Widespread Transcriptomic Changes in CD4+ T Cells Early Postinfection. mBio. 12(3):e01369–21.

65. Greenwood EJD, Williamson JC, Sienkiewicz A, Naamati A, Matheson NJ, Lehner PJ. Promiscuous Targeting of Cellular Proteins by Vpr Drives Systems-Level Proteomic Remodeling in HIV-1 Infection. Cell Reports. 2019 Apr 30;27(5):1579–1596.e7.

66. Reuschl AK, Mesner D, Shivkumar M, Whelan MVX, Pallett LJ, Guerra-Assunção JA, et al. HIV-1 Vpr drives a tissue residency-like phenotype during selective infection of resting memory T cells. Cell Reports. 2022 Apr 12;39(2):110650.

67. Badley AD, Sainski A, Wightman F, Lewin SR. Altering cell death pathways as an approach to cure HIV infection. Cell Death Dis. 2013 Jul 11;4(7):e718.

68. Cummins NW, Sainski AM, Dai H, Natesampillai S, Pang YP, Bren GD, et al. Prime, Shock, and Kill: Priming CD4 T Cells from HIV Patients with a BCL-2 Antagonist before HIV Reactivation Reduces HIV Reservoir Size. Journal of Virology [Internet]. 2016 Feb 3 [cited 2024 Dec 5]; Available from: https://journals.asm.org/doi/10.1128/jvi.03179-15

69. Archin NM, Eron JJ, Palmer S, Hartmann-Duff A, Martinson JA, Wiegand A, et al. Valproic acid without intensified antiviral therapy has limited impact on persistent HIV infection of resting CD4+ T cells. AIDS. 2008 Jun 19;22(10):1131.

70. Spivak AM, Andrade A, Eisele E, Hoh R, Bacchetti P, Bumpus NN, et al. A Pilot Study Assessing the Safety and Latency-Reversing Activity of Disulfiram in HIV-1–Infected Adults on Antiretroviral Therapy. Clinical Infectious Diseases. 2014 Mar 15;58(6):883–90.

71. Cillo AR, Sobolewski MD, Bosch RJ, Fyne E, Piatak M, Coffin JM, et al. Quantification of HIV-1 latency reversal in resting CD4+ T cells from patients on suppressive antiretroviral therapy. Proceedings of the National Academy of Sciences. 2014 May 13;111(19):7078–83.

72. Busca A, Saxena M, Kumar A. Critical Role for Antiapoptotic Bcl-xL and Mcl-1 in Human Macrophage Survival and Cellular IAP1/2 (cIAP1/2) in Resistance to HIV-Vpr-induced Apoptosis. J Biol Chem. 2012 Apr 27;287(18):15118–33.

73. Goonetilleke N, Clutton G, Swanstrom R, Joseph SB. Blocking Formation of the Stable HIV Reservoir: A New Perspective for HIV-1 Cure. Frontiers in Immunology [Internet]. 2019 [cited 2023 Apr 4];10. Available from: https://www.frontiersin.org/articles/10.3389/fimmu.2019.01966

74. Wen X, Duus KM, Friedrich TD, Noronha CMC de. The HIV1 Protein Vpr Acts to Promote G2 Cell Cycle Arrest by Engaging a DDB1 and Cullin4A-containing Ubiquitin Ligase Complex Using VprBP/DCAF1 as an Adaptor *. Journal of Biological Chemistry. 2007 Sep 14;282(37):27046–57.

75. Love MI, Huber W, Anders S. Moderated estimation of fold change and dispersion for RNA-seq data with DESeq2. Genome Biology. 2014 Dec 5;15(12):550.

